# Metabolite profiling of experimental cutaneous leishmaniasis lesions demonstrates significant perturbations in tissue phospholipids

**DOI:** 10.1101/2020.05.13.094649

**Authors:** Adwaita R. Parab, Diane Thomas, Sharon Lostracco-Johnson, Jair Lage de Siqueira-Neto, James McKerrow, Pieter C. Dorrestein, Laura-Isobel McCall

**Affiliations:** Department of Microbiology and Plant Biology, University of Oklahoma, Norman, Oklahoma, United States of America; Laboratories of Molecular Anthropology and Microbiome Research, University of Oklahoma, Norman, Oklahoma, United States of America; Skaggs School of Pharmacy and Pharmaceutical Sciences, University of California San Diego, La Jolla, California, United States of America; Center for Microbiome Innovation, University of California San Diego, La Jolla, California, United States of America; Collaborative Mass Spectrometry Innovation Center, University of California San Diego, La Jolla, California, United States of America; Department of Chemistry and Biochemistry, University of Oklahoma, Norman, Oklahoma, United States of America

## Abstract

Each year 700,000 to 1.2 million new cases of cutaneous leishmaniasis (CL) are reported and yet CL remains one of thirteen diseases classified as neglected tropical diseases (NTDs). *Leishmania major* is one of several different species of that same genus that can cause CL. Current CL treatments are limited by adverse effects and rising resistance. Studying disease metabolism at the site of infection can lead to new drug targets. In this study, samples were collected from mice infected in the ear and footpad with *L. major* and analyzed by untargeted liquid chromatography-tandem mass spectrometry (LC-MS/MS). Significant differences in overall metabolite profiles were noted in the ear at the site of the lesion. Interestingly, lesion-adjacent, macroscopically healthy sites also showed alterations in specific metabolites, including select phosphocholines (PCs). Host-derived PCs in the lower *m/z* range (*m/z* 200-799) showed an increase with infection in the ear at the lesion site, while those in the higher *m/z* range (*m/z* 800-899) were decreased with infection at the lesion site. Overall, our results expanded our understanding of the mechanisms of CL pathogenesis through the host metabolism and may lead to new curative measures against infection with *Leishmania*.

**Author summary:** Cutaneous leishmaniasis (CL) is one of thirteen neglected tropical diseases in the world today. It is an infectious disease with a wide distribution spanning five continents, with increasing distribution expected due to climate change. CL manifests as skin lesions and ulcers that are disabling and stigmatized. With the current treatment options being limited, studying host-pathogen metabolism can uncover mechanisms of disease pathogenesis that may lead to new curative measures against infection. In this paper we used untargeted metabolomics to address molecular-level changes occurring *in vivo* in experimental skin lesions of *Leishmania major*. Distinct global metabolic profiles were observed. Total phosphocholines (PCs) and those in the lower *m/z* ranges were significantly higher at the site of the skin lesion in the ear. In addition, specific PCs as well as PCs of varied *m/z* ranges were also affected at healthy-appearing lesion-adjacent sites, indicating that infection-induced metabolic perturbations are not restricted to the lesion site. Ultimately, these results provide essential clues to the metabolic pathways affected by CL.

## Introduction

Leishmaniasis affects people in 88 countries worldwide in tropical, subtropical and temperate regions, putting approximately 350 million individuals at risk of infection, with approximately 12 million battling the disease [1]. It is one of the three most impactful vector-borne protozoan neglected tropical diseases, causing approximately 2.1 million DALYs (Disability-Adjusted Life Years) and 51,000 deaths. With recent population movements, leishmaniasis is now affecting people in non-endemic regions as well. The expanding spread of leishmaniasis can be attributed to climate change and social constraints of populations living in poverty and conflict. Leishmaniasis is a disease that is exacerbated by poverty and socio-economic barriers, increasing rates of disease progression, mortality and morbidity [2]. It comes at the high cost of treatment with the consequences of low or no income due to social stigmas associated with the symptoms of skin lesions, ulcers and disfigurement. Ultimately it puts financial burdens on individuals as well as societies as a whole [3].

Leishmaniasis is caused by about 20 different species of the parasite *Leishmania,* with three clinical syndromes: visceral, cutaneous (CL) and mucocutaneous leishmaniasis. CL is the most common form of the disease and symptoms include skin lesions and ulcers on exposed parts of the body. Mucocutaneous leishmaniasis is a disabling form in which the lesions can lead to destruction of soft tissue of the nose, mouth and throat cavities. Of the three clinical forms of the disease, visceral leishmaniasis (kala-azar) is the most deadly, with serious symptoms like swelling of the liver and spleen, extreme anemia and frequent bouts of fever. Infection is transmitted through female sand-flies of the *Phlebotomus* genus in the Old World and *Lutzomyia* genus in the New World [4]. Promastigotes enter the body upon being bitten by a female sandfly. They are taken up by macrophages, where they enter the amastigote stage, multiplying and affecting various tissue types depending on whether infection is initiated by a viscerotropic or dermotropic parasite strain [5][6]. This initiates the clinical manifestations of the disease. Humans as well as other mammals serve as host reservoirs for the parasite [7].

The current course of treatment for CL is usually antimonial drug compounds. These are known to be highly toxic compounds, in addition to the threat of increased parasite-resistance to antimony in several regions of the world. Miltefosine, amphotericin B and paromomycin are among the other drugs that are administered for CL treatment, all of which have the drawbacks of high level of toxicity, increased drug resistance and treatment failure. Miltefosine is also teratogenic and should not be given to women in childbearing age. Treatment failure can be attributed to the characteristics of the host (immune system and nutritional status), of the parasite (mechanisms of survival within the host, drug resistance mechanisms, tissue location, etc.) and environmental factors such as awareness and treatment accessibility [8]. Approaching disease pathogenesis from a molecular perspective could uncover new mechanisms of infection and aid in developing new cures for leishmaniasis [9].

Alongside genes and proteins, metabolites play an important role in the life of an organism. The metabolome reflects the true functional endpoint of a complex biological system and provides a functional view of the organism by taking into account the sum of its genes, RNA, proteins and its environment [10]. Untargeted metabolomics can help identify metabolites involved in disease pathogenesis, in an unbiased fashion, acquiring data across a broad mass range [11]. For example, untargeted metabolomics has shown that miltefosine’s mode of action may be related to modulation of parasite lipid metabolism, particularly increased levels of by-products of lipid turnover [12]. The overall aim of this work was to perform an untargeted metabolic analysis of CL lesions in mice infected with *Leishmania major*. Our results showed significant changes in the host metabolism, specifically the PC pathway, in the skin lesions of CL.

## Methods

### Ethics statement

All vertebrate animal studies were performed in accordance with the USDA Animal Welfare Act and the Guide for the Care and Use of Laboratory Animals of the National Institutes of Health. Euthanasia was performed by isoflurane overdose followed by cervical dislocation, under a protocol approved by the University of California San Diego Institutional Animal Care and Use Committee (protocol S14187).

### *In vivo* experimentation

Female BALB/c mice (6-8 week-old) were injected intradermally in the left ear with 1×10^6^ luciferase-expressing *L. major* strain LV39 promastigotes or in the left rear footpad with 5×10^6^ luciferase-expressing *L. major* strain LV39 promastigotes in PBS [13]. Infected and uninfected ear tissue were collected 8 weeks post-infection and infected and uninfected footpads were collected 7 weeks post-infection, and immediately snap-frozen. Samples were stored at - 80°C until metabolite extraction. Parasites were maintained at 28 °C in M199 medium (Sigma) supplemented with 10% fetal bovine serum (Sigma), 1% penicillin-streptomycin, RPMI 1640 vitamin mix (1%), HEPES (25 mM), adenosine (100 μM), glutamine (1 mM), hemin (0.005%), NaHCO3 (12 mM) and folic acid (10 μM) (pH 7.2) [14].

### LC-MS/MS

Metabolite extraction, liquid chromatography and mass spectrometry were performed as previously described [15]. Briefly, metabolites were extracted with 50% methanol (aqueous extract) followed by 3:1 dichloromethane:methanol (organic extract). LC was performed on an UltiMate 3000 UHPLC (Thermo Scientific) with Phenomenex UHPLC 1.7 µm 100 Å Kinetex C8 column (50 × 2.1 mm), and with water and 0.1% formic acid as mobile phase A and acetonitrile and 0.1% formic acid as mobile phase B, flow rate of 0.5 mL/min and column temperature of 40°C. Daily MS calibration was performed with ESI-L Low Concentration Tuning Mix (Agilent Technologies). The internal calibrant was Hexakis(1H,1H,3H-tetrafluoropropoxy)phosphazene (Synquest Laboratories), *m/z* 922.009798 which was present throughout the run. MS/MS data for each run was collected by fragmentation of the ten most intense ions, in range 80-2,000 *m/z*, with active exclusion after 4 spectra and release after 30s. LC gradients and MS parameters for each extraction were as follows (Table 1 and Table 2).

**Table 1.** LC Gradients.

| Ear aqueous extraction |  |
| --- | --- |
| Start | 2% B |
| 1 min | 2% B |
| 1.5 min | 40% B |
| 4 min | 98% B |
| 5 min | 98% B |
| 6 min | 2% B |
| 7 min | 2% B |
| **Ear organic extraction** |  |
| Start | 2% B |
| 1 min | 2% B |
| 1.5 min | 60% B |
| 5.5 min | 98% B |
| 7.5 min | 98% B |
| 8.5 min | 2% B |
| 10.5 min | 2% B |
| **Footpad aqueous extraction** |  |
| Start | 2% B |
| 1 min | 2% B |
| 1.5 min | 40% B |
| 6 min | 98% B |
| 6.5 min | 98% B |
| 7 min | 2% B |
| **Footpad organic extraction** |  |
| Start | 2% B |
| 1 min | 2% B |
| 1.5 min | 70% B |
| 7 min | 98% B |
| 8 min | 98% B |
| 9 min | 2% B |
| 10.5 min | 2% B |

**Table 2.**
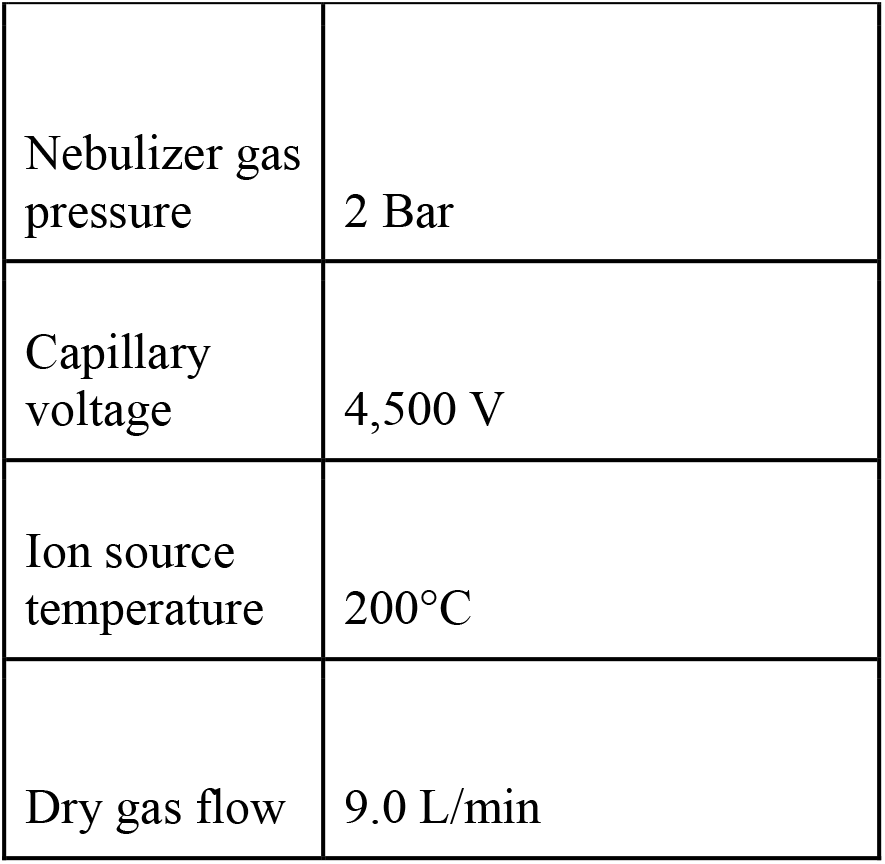
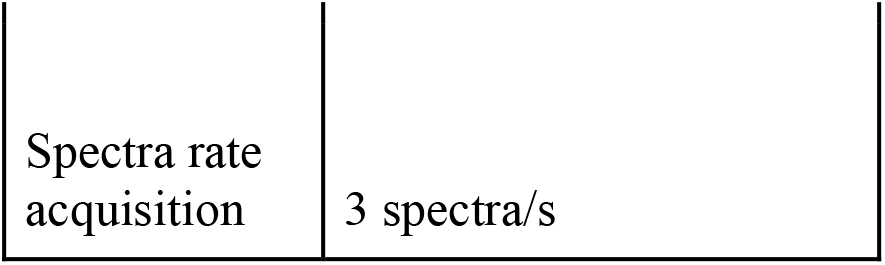
MS parameters.

|  |  |
| --- | --- |
| Nebulizer gas pressure | 2 Bar |
| Capillary voltage | 4,500 V |
| Ion source temperature | 200°C |
| Dry gas flow | 9.0 L/min |
| Spectra rate acquisition | 3 spectra/s |

### LC-MS/MS Data analysis

LC-MS/MS data was processed using MZmine 2.37 [16], with parameters as shown in Table 3.

**Table 3.** MZmine parameters.

| Mass Detection |  |
| --- | --- |
| MS level 1: Noise level | 1.00E+03 |
| MS level 2: Noise level | 10 |
| Mass detector | Centroid |
| Chromatogram Builder |  |
| Min time span | 0.06 min |
| Min peak height | 3.00E+03 |
| *m/z* tolerance | 1e-6 or 10 ppm |
| Chromatogram Deconvolution |  |
| Algorithm | Baseline cutoff |
| Min peak height | 3.00E+03 |
| Peak duration range (min) | 0.06-2 min (ear), 0.01-7 min (footpad) |
| Baseline level | 1.00E+02 (ear), 1.50E+03 (footpad) |
| *m/z* range for MS2 scan pairing (Da) | 0.01 |
| RT range for MS2 scan pairing (min) | 0.2 min |
| **Isotopic Peaks Grouper** |  |
| *m/z* tolerance | 1e-6 or 10 ppm |
| Retention time tolerance (absolute: min) | 0.05 min |
| Monotonic shape | Enabled |
| Maximum charge | 3 |
| Representative isotope | Most intense |
| **Join Aligner** |  |
| *m/z* tolerance | 1e-6 or 10 ppm |
| Weight for *m/z* | 7 |
| Retention time tolerance (absolute: min) | 0.5 min |
| Weight for RT | 3 |
| **Manual Filtering** |  |
| Min number of peaks per row | 3 |
| RT range | 0.2-10.5 (ear organic and footpad),<br>0.2-6.9 (ear aqueous), |
| MS2 | required |
| Manual validation of peak shape |  |
| **Gap-filing** |  |
| *m/z* tolerance | 0.000001 or 10 ppm |
| RT tolerance | 0.5 min |
| Intensity tolerance | 30% |
| RT correction | Enabled |

Total ion current (TIC) normalization and data processing was performed in Jupyter notebook with R [17]. Principal Coordinate Analysis (PCoA) was done using the Bray-Curtis-Faith dissimilarity matrix implemented in QIIME1 [18] and PERMANOVA calculations were performed using the R package “vegan” to compare the chemical similarity of samples from the four groups of varying condition and position of infection [19,20]. EMPeror was used to visualize PCoA plots [21]. randomForest package in R was used to find variables of importance associated with infection and sampling conditions, using 7000 trees [22]. Global Natural Products Social Molecular Networking platform (GNPS) was used to annotate molecules from spectral library references and to perform feature-based molecular networking [23][24][25]. The following parameters were used in GNPS: precursor ion mass tolerance of 0.02 Da, fragment ion mass tolerance of 0.02 Da, minimum cosine score of 0.7 and 4 or more matched fragment ions. The maximum shift allowed between two MS/MS spectra was 500 Da, 10 maximum neighbor nodes allowed and maximum difference between precursor ion mass of searched MS/MS spectrum and library spectra was 100 Da. Spectral matches were evaluated by considering cosine scores, quality of mirror plots, as well as the number of matched peaks. Molecular network visualization was done in Cytoscape 3.7.2 [26]. Notched box plots showing metabolite feature abundance for the four different groups (infected/uninfected vs. center/edge) for the ear samples and two different groups (infected vs. uninfected) for the footpad samples along with non-parametric two-tailed Wilcoxon statistical tests were both performed in R. Boxplot whiskers represent the lowest and largest data points and non-overlapping boxplot notches indicate different medians between groups (95% confidence).

### Data Availability Statement

Data has been deposited in MassIVE (massive.ucsd.edu, accession numbers MSV000081004 (ear) and MSV000080239 (footpad)). Molecular networking can be accessed here: https://gnps.ucsd.edu/ProteoSAFe/status.jsp?task=451754c383de461e9e4abdf6eb3199d2 (aqueous ear extraction), https://gnps.ucsd.edu/ProteoSAFe/status.jsp?task=0d092bbb213347c3bd7a19b9cae2bcf4 (organic ear extraction), https://gnps.ucsd.edu/ProteoSAFe/status.jsp?task=becfa09afe7b4f83a7c5621029f2df24 (aqueous footpad extraction), https://gnps.ucsd.edu/ProteoSAFe/status.jsp?task=fb6f32dcafe34ec587bb264341814217 (organic footpad extraction).

## Results

To better understand the impact of infection on tissue metabolites, we analyzed overall and specific metabolite differences in the presence and absence of infection with *L. major,* at sites of lesion and lesion-adjacent sites (with no visible signs of infection). BALB/c mice were injected intradermally in the ear. Eight weeks post-infection, samples were collected from the area where the parasites were injected, which showed skin lesions (“infected ear center”), the surrounding area that appeared infection-free (“infected ear edge”), and the matched tissue regions from the uninfected ear (“uninfected ear center”, “uninfected ear edge”) (Fig 1 A). Metabolites were extracted with aqueous and organic solvents and analyzed by untargeted LC-MS/MS (see Methods). Overall, for both aqueous and organic extractions, distinct global metabolite profiles were observed by Principal Coordinate Analysis (PCoA) for the infected ear center compared to infected ear edge (PERMANOVA p<0.01, aqueous extraction R^2^ = 0.743, organic extraction R^2^= 0.643, Fig 1 B and 1 C), to uninfected ear center (PERMANOVA p<0.01, aqueous extraction R^2^=0.739, organic extraction R^2^=0.805, Fig 1 B and 1 C) and to uninfected ear edge (PERMANOVA p<0.01, aqueous extraction R^2^=0.288, organic extraction R^2^=0.248, Fig 1 B and 1 C). In contrast, no significant differences for both aqueous and organic extracts by PCoA analysis in terms of overall metabolite profile were observed between infected ear edge and uninfected ear samples (PERMANOVA p>0.1, Fig 1 B and 1 C). Thus, *L. major* infection changes the overall tissue chemical composition at the lesion location in the ear. In contrast, the impact of *L. major* infection on overall footpad metabolite profile for the organic (PERMANOVA p=0.218 R^2^=0.156, S1 Fig) and aqueous (PERMANOVA p=0.244 R^2^=0.146, S1 Fig) extractions was much more minor.

**Fig.1.**
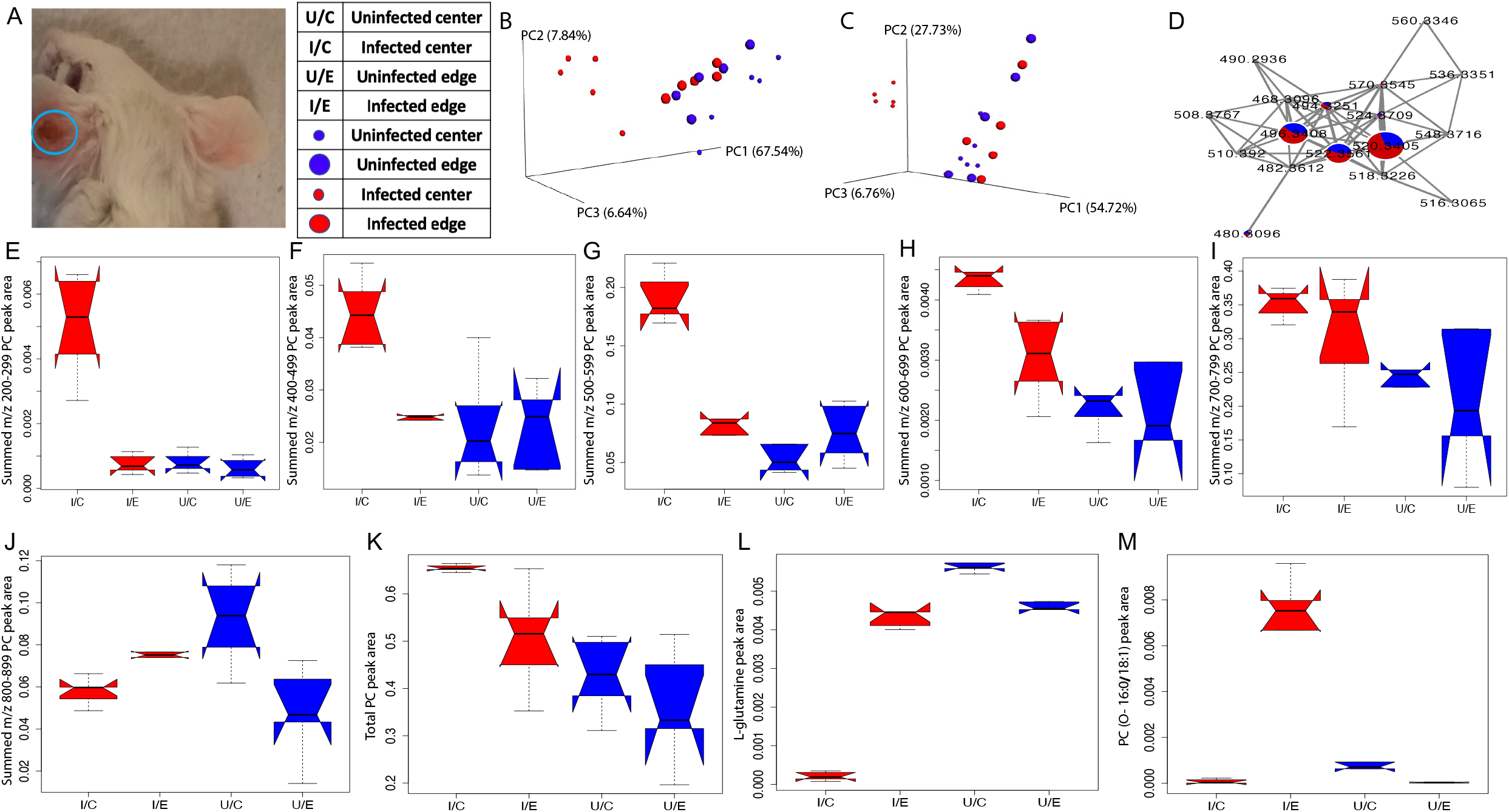
Effect of *in vivo L. major* infection on host metabolite profile. (A) Sites of infection and sample collection. Lesion at the center of the infected ear is circled in blue. (B) PCoA analysis of aqueous extraction from infected and uninfected ear samples, showing overall differences in metabolite profiles between sampling sites. PERMANOVA p= 0.004, R^2^=0.288. (C) PCoA analysis of organic extraction from infected and uninfected ears, showing differences in global metabolite profiles between sampling sites. PERMANOVA p= 0.003, R^2^=0.248. (D) Representative subnetwork of phosphocholine (PC) molecular family members found in ear tissue and showing high relative abundance with infection (red) and low abundance without infection (blue). (E-J) PCs in the *m/z* range 200-299, 400-499, 500-599, 600-699, 700-799 and 800-899, respectively, change with infection and sampling position in the ear. (K) Total PC levels were increased at the site of infection in the ear. Non-overlapping boxplot notches indicate significantly different medians between groups [27]. (L) Representative metabolite decreased by infection at the site of the lesion: glutamine (Wilcoxon rank-sum test comparing infected ear center vs infected ear edge p=0.007937). (M) Representative metabolite increased only at infection-adjacent sites: 1-hexadecyl-2-(9Z-octadecenoyl)-sn-glycero-3-phosphocholine (PC(O-16:0/18:1), Wilcoxon rank-sum test comparing infected ear center vs infected ear edge p=0.007937).

Random forest machine learning analysis [22] was performed to identify the metabolites most affected by infection in both experimental systems, with annotation performed using molecular networking and GNPS [23]. Annotatable molecules most highly affected by infection include metabolites of the phosphocholine (PC) family of phospholipids, glutamine, and eicosatrienoic acid (Table 4, 5, 6, 7, S2 Fig). Glutamine was decreased with infection at the site of the ear lesion (Wilcoxon rank sum test p value=0.0079 compared to the uninfected ear center, Fig 1 L), although it was unaffected by infection in the footpad, and eicosatrienoic acid was increased in the infected footpad (Wilcoxon rank sum test p value=0.0079). Given that many of the differential molecules are PCs, we investigated the impact of infection on this family in greater detail. Molecular network analysis of PC family molecules in both aqueous and organic ear extracts showed that most detected PCs were strongly affected by infection (Fig 1 D, S3 Fig). In particular, total PCs and PCs in the lower ranges of *m/z* 200-299, 400-499, 500-599, 600-699 were significantly higher in the infected ear center compared to the infected ear edge, to uninfected ear center, and to uninfected ear edge, as well as for all uninfected samples (both positions combined) vs all infected samples (both positions combined) (Wilcoxon rank sum test p value<0.05 for each pairwise comparison, Fig 1 E, F, G, H, K). Given that all these PCs were detected in both infected and uninfected samples, albeit at differential abundances, they are host-derived rather than parasite-derived. No PCs were detected in the *m/z* 300-399 range. PCs in the range of *m/z* 700-799 showed a similar trend where the infected and uninfected sample groups were significantly different (Wilcoxon rank sum test p value<0.05). PCs in the range of *m/z* 700-799 were significantly higher in the infected ear center compared to the uninfected ear edge and center (Wilcoxon rank sum test p value<0.05, Fig 1 I), while the levels of PCs were comparable between infected ear center and edge (Wilcoxon rank sum test p value = 0.55, Fig 1 I). In the *m/z* 800-899 range the opposite trend was seen, where the uninfected ear samples were not significantly different from the infected ear (Wilcoxon sum rank test p value=0.9, Fig 1 J). However, PCs in the range *m/z* 800-899 were significantly higher in the uninfected ear center than the infected ear center (Wilcoxon rank sum test p value<0.05, Fig 1 J). In contrast, the footpad PCs aggregated into *m/z* ranges did not show significant differences between the infected and uninfected groups, although specific PCs were increased by infection in the footpad (Table 6, 7). These results indicate that PCs are strongly affected by cutaneous *Leishmania* infection. In addition, our observation that specific PCs as well as PCs of varied *m/z* ranges are also affected at lesion-adjacent sites (“infected ear edge”) indicates that infection-induced metabolic perturbations are not restricted to the lesion site, revealing a better picture of what is happening to the host during the disease state and providing clues to the pathways involved.

**Table 4.** Most differential metabolite features (as identified by random forest analysis) for ear aqueous extraction.

| *m/z* | RT | Spectral match on GNPS / LIPID MAPS annotation | Mass difference | PPM error | Cosine score | Number of matched peaks | Impact of infection in the ear | P value (infected ear center vs uninfected ear center) | P value (infected ear edge vs uninfected ear edge) | P values (infected ear vs uninfected ear) | Effect in Footpad (Aqueous extraction) | Effect in Footpad (Organic extraction) |
| --- | --- | --- | --- | --- | --- | --- | --- | --- | --- | --- | --- | --- |
| 794.6051 | 4.42 | PC family member; LipidMAPS: PC(O-38:5) | NA | NA | NA | NA | increased in infected ear edge | 0.056 | 0.0079 | 0.48 | ND | increased in infected footpad |
| 772.6201 | 4.73 | PC family member | NA | NA | NA | NA | increased in infected ear edge | 0.016 | 0.0079 | 0.39 | ND | ND |
| 750.5435 | 5.06 | PC family member | NA | NA | NA | NA | increased in infected ear center | 0.0079 | 0.0079 | 2.2E-05 | ND | unaffected by infection |
| 155.0498 | 0.85 | NA | NA | NA | NA | NA | increased in infected & uninfected ear edges | 0.0079 | 0.22 | 0.68 | increased in infected footpads | ND |
| 752.5585 | 5.07 | NA | NA | NA | NA | NA | increased in infected ear center | 0.0079 | 0.0079 | 1.1E-05 | ND | increased in infected footpad (not statistically significant) |
| 744.5906 | 4.48 | PC family member | NA | NA | NA | NA | increased in infected ear edge | 0.0079 | 0.0079 | 0.052 | ND | increased in infected footpad (not statistically significant) |
| 746.6052 | 4.78 | 1-headecyl-2-(9Z-octadecenoyl)-sn-glycero-3-phosphocholine | 0 | 4 | 0.96 | 14 | increased in infected ear edge | 0.032 | 0.0079 | 0.28 | ND | increased in infected footpad |
| 796.6205 | 4.91 | PC family member | NA | NA | NA | NA | increased in infected ear edge | 0.15 | 0.0079 | 0.11 | ND | increased in infected footpad |
| 147.0815 | 0.34 | glutamine | 0.005 | 31 | 0.87 | 4 | decreased in infected ear center | 0.0079 | 0.42 | 0.00073 | unaffected by infection | unaffected by infection |
| 770.605 | 4.58 | PC family member | NA | NA | NA | NA | increased in infected ear edge | 0.31 | 0.0079 | 0.22 | ND | unaffected by infection |
| 720.5887 | 4.7 | PC family member | NA | NA | NA | NA | increased in infected ear edge | 0.095 | 0.0079 | 0.28 | ND | increased in infected footpad |
| 794.6052 | 6.13 | PC(O-38:5) | NA | NA | NA | NA | increased in infected ear center | 0.0079 | 0.0079 | 1.1E-05 | ND | increased in infected footpad |
| 330.1314 | 6.4 | NA | NA | NA | NA | NA | increased in infected<br>ear center | 0.0079 | 0.0079 | 1.1E-05 | ND | ND |
| 169.0624 | 0.31 | NA | NA | NA | NA | NA | decreased<br>in<br>infected<br>ear center | 0.011 | 0.55 | 0.011 | unaffected<br>by<br>infection | unaffected<br>by<br>infection |
| 261.1474 | 0.47 | NA | NA | NA | NA | NA | increased<br>in<br>infected<br>&<br>uninfected<br>ear edges | 0.095 | 0.31 | 0.31 | unaffected<br>by<br>infection | unaffected<br>by<br>infection |

NA: not applicable; ND: not detected

**Table 5.**
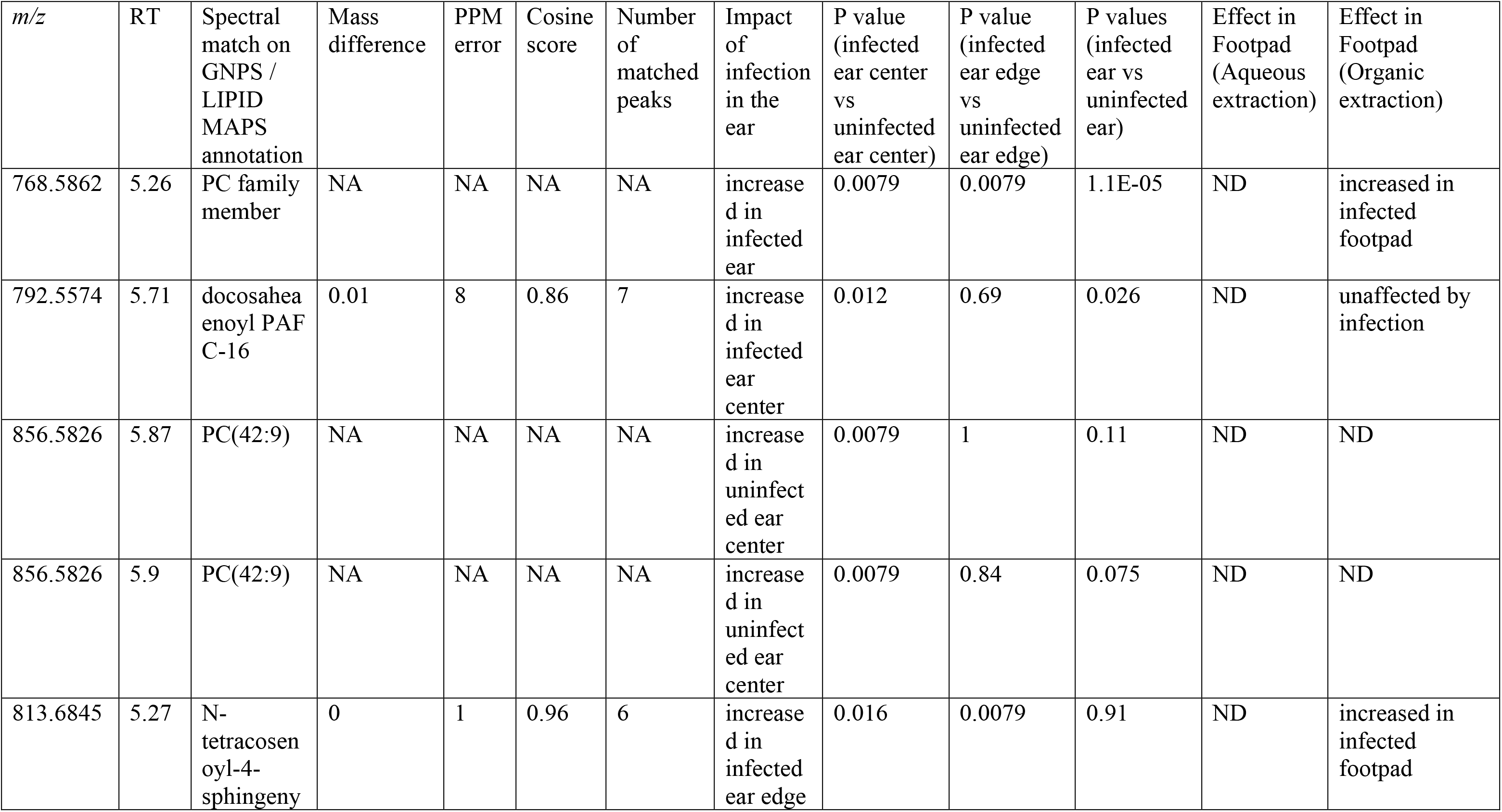
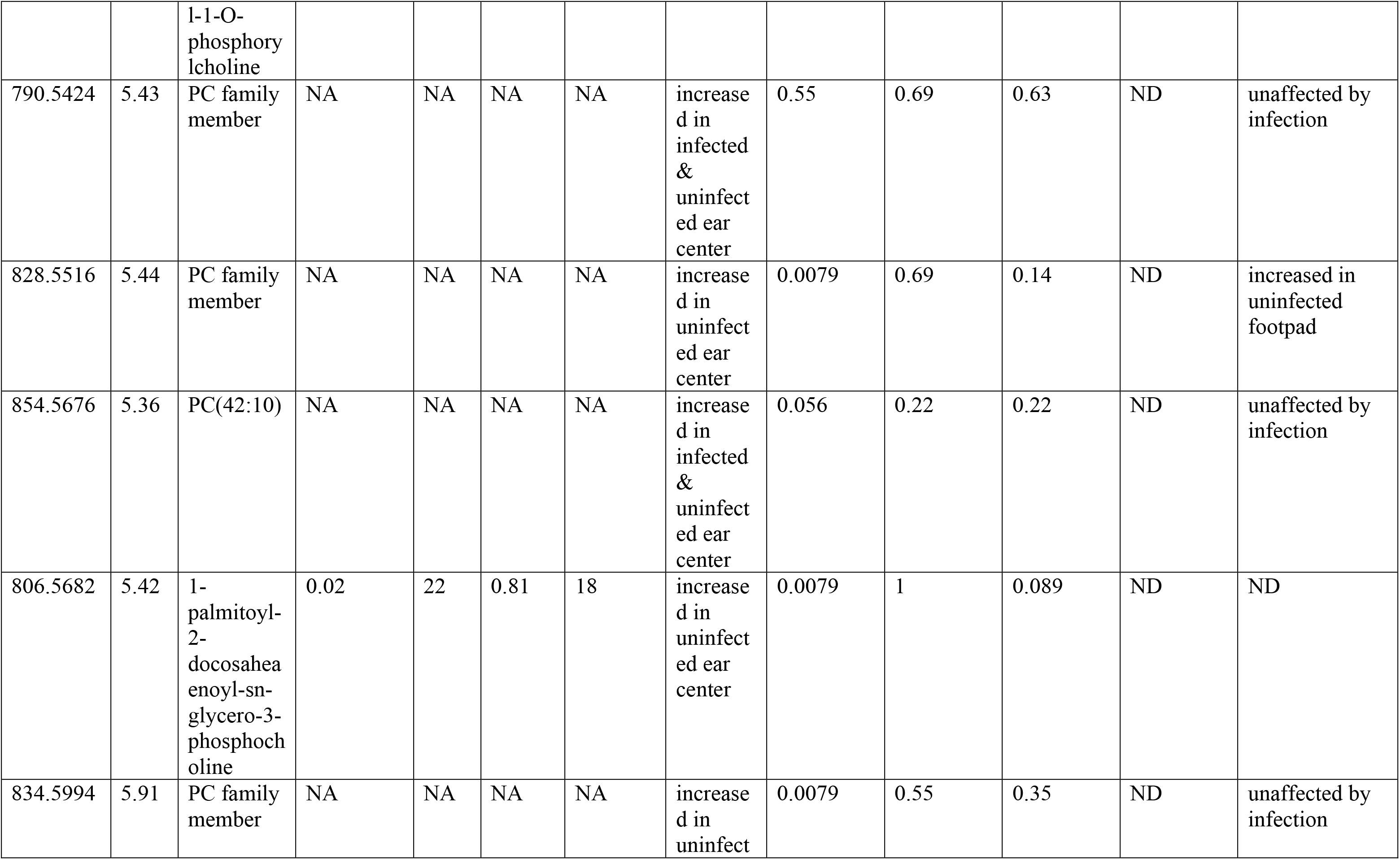
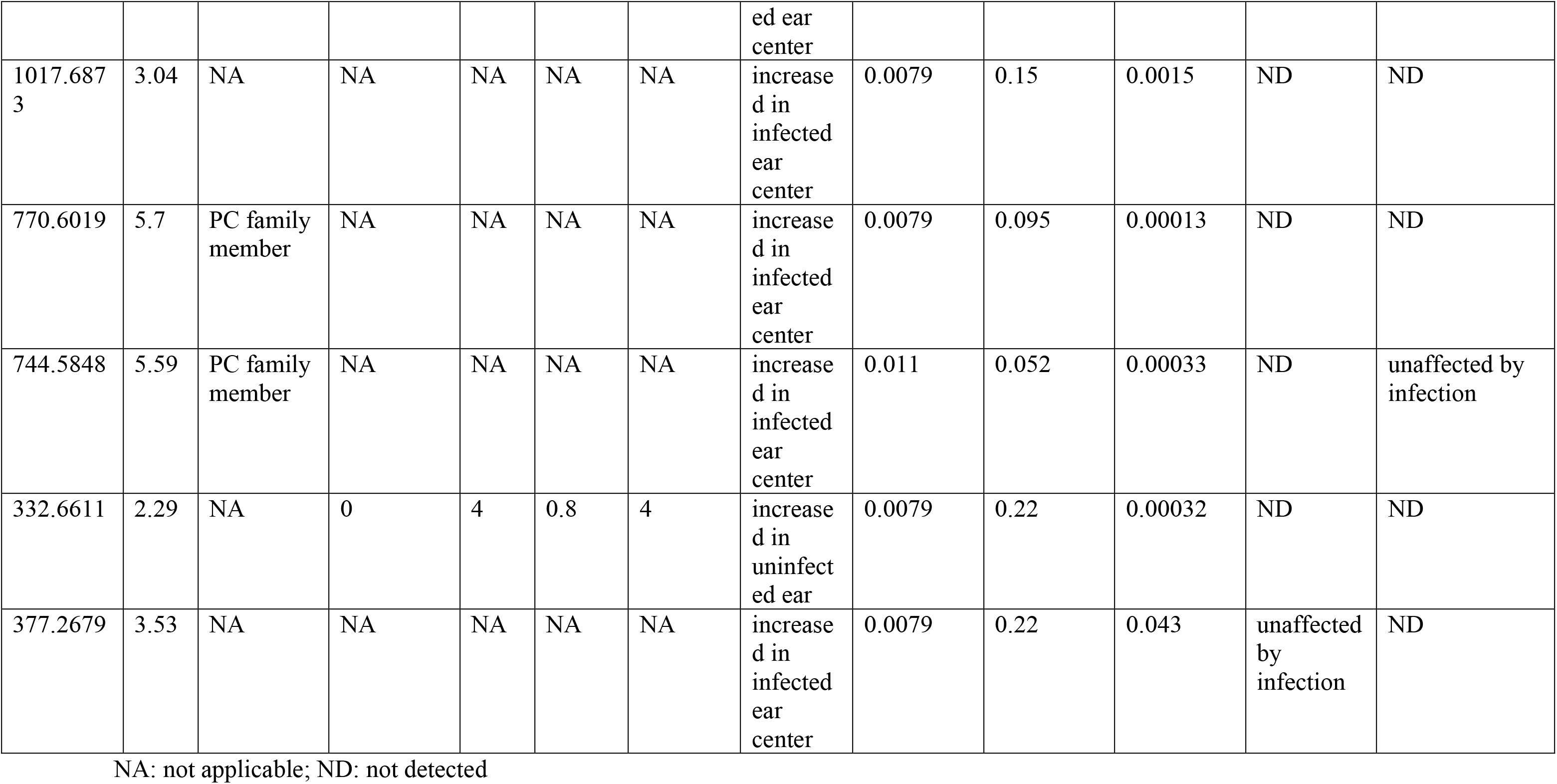
Most differential metabolite features (as identified by random forest analysis) for ear organic extraction.

**Table 6.** Most differential metabolite features (as identified by random forest analysis) for footpad aqueous extraction.

| *m/z* | RT | Spectral match on GNPS / LIPID MAPS annotation | Mass difference | PPM error | Cosine score | Number of matched peaks | Impact of infection | P values |
| --- | --- | --- | --- | --- | --- | --- | --- | --- |
| 331.2638 | 4.31 | NA | NA | NA | NA | NA | increased in infected footpad | 0.0079 |
| 368.2591 | 4.06 | acylcarnitine family member | NA | NA | NA | NA | increased in infected footpad | 0.0079 |
| 377.1461 | 2.41 | NA | NA | NA | NA | NA | increased in infected footpad | 0.012 |
| 425.3375 | 3.18 | NA | NA | NA | NA | NA | increased in uninfected footpad | 0.0079 |
| 210.1121 | 2.74 | NA | NA | NA | NA | NA | increased in infected footpad | 0.0079 |
| 212.1651 | 2.75 | NA | NA | NA | NA | NA | increased in uninfected footpad | 0.0079 |
| 206.1067 | 4.51 | NA | NA | NA | NA | NA | increased in infected footpad | 0.0079 |
| 549.2233 | 2.49 | NA | NA | NA | NA | NA | increased in uninfected footpad | 0.0079 |
| 522.2834 | 4.16 | 1-(9Z-octadecenoyl)-sn-glycero-3-phosphocholine | 0 | 0 | 0.95 | 15 | increased in infected footpad | 0.0079 |
| 303.2323 | 4.1 | 5,6-epoy-8Z,11Z,14Z-eicosatrienoic acid | 0 | 4 | 0.89 | 8 | increased in infected footpad | 0.0079 |
| 327.2325 | 4.05 | NA |  |  |  |  | increased in infected footpad | 0.0079 |
| 508.3764 | 4.01 | 1-(1Z-octadecenyl)-sn-glycero-3-phosphocholine | 0 | 2 | 0.91 | 10 | increased in infected footpad | 0.0079 |
| 281.0052 | 2.55 | NA | NA | NA | NA | NA | increased in<br>infected footpad | 0.0079 |
| 377.2661 | 4.32 | NA | NA | NA | NA | NA | increased in<br>infected footpad | 0.0079 |
| 230.1756 | 2.74 | NA | NA | NA | NA | NA | increased in<br>uninfected<br>footpad | 0.0079 |

NA: not applicable

**Table 7.**
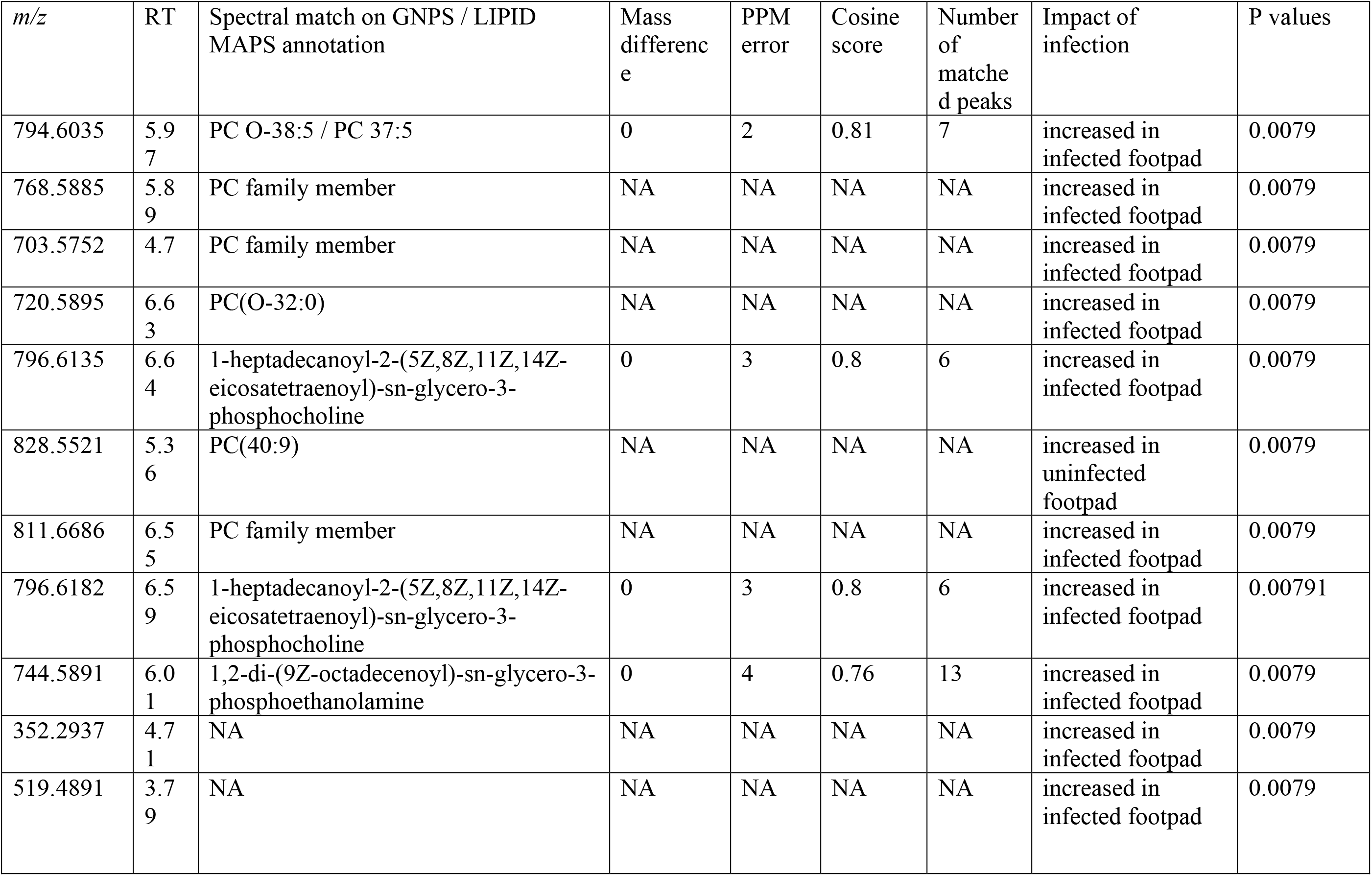
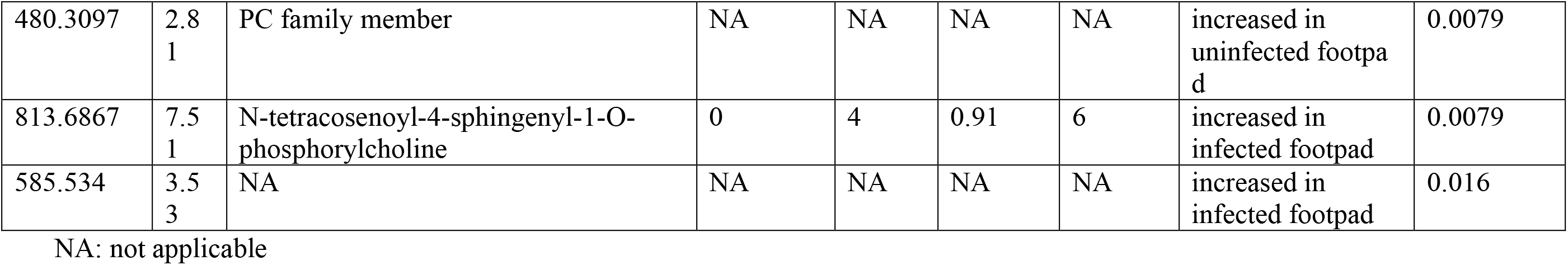
Most differential metabolite features (as identified by random forest analysis) for footpad organic extraction.

## Discussion

The metabolome provides a link between genotype and phenotype by identifying changes occurring at the molecular level, for example when parasites and their hosts interact [28]. Metabolism is also an indicator of the host physiological state. Understanding the infection-induced host metabolic alterations could lead to new treatments for parasitic diseases [29], particularly host-targeted drug therapy focused on pathways otherwise redundant to the host but important for parasite invasion, replication and survival [30], or on mitigating damage caused by the parasite [29]. In addition, host metabolism, as measured in plasma samples, has been shown to be able to serve as an indicator of response to CL treatment [31]. Several studies have investigated *Leishmania* metabolism during *in vitro* macrophage infection (*e.g.* [32]), or in amastigotes purified from mouse granulomatous lesions [33], but there is still a lack of knowledge of host metabolic responses during *in vivo* infection. Given the relative host vs parasite biomass and the slow replication of *Leishmania* during *in vivo* infection [33], it is likely that most metabolites identified in our study were host-derived, thereby expanding our understanding of host metabolic contributions to CL pathogenesis.

Amongst annotatable metabolites in our study, members of the PC family were most affected by infection. PCs were detected in both infected and uninfected groups. PCs of the smaller *m/z* range (*m/z* 200-799) were significantly higher with infection and those in the larger *m/z* range (*m/z* 800-899) showed the opposite trend, where PC levels were decreased by infection at the lesion site (Fig. 1E-J). Total PCs were significantly higher in the infected ear center vs the infected ear edge, uninfected ear center and uninfected ear edge (Fig. 1K). Select PCs were also increased in the infected footpad (Tables 6-7). Miltefosine is a commonly administered oral drug for the treatment of visceral and CL that targets the PC biosynthetic pathway [34]. Importantly, miltefosine was originally developed for its anti-tumor properties against cancer, and as such can be expected to also proceed via host-directed effects in addition to impacts on parasite metabolism. We speculate that miltefosine mechanism of action in CL may thus also involve re-normalization of infection-induced changes in host PCs. Future studies are thus needed to investigate the mechanism of action of miltefosine with respect to host metabolism in CL.

Additional annotatable metabolites also included the omega-3 fatty acid eicosatrienoic acid and glutamine. Glutamine was noted to be significantly lower with infection at the site of the ear lesion. A recent study in mice infected with *L. donovani* showed heightened glutamine consumption with infection and a role of glutamine supplementation in clearing parasite load [35]. Additionally, glutamine uptake is also essential to the pathogenesis of *Toxoplasmosis gondii* parasite infection [36]. Future studies should aim to look at the specific functional role of glutamine metabolism in *L. major* infection.

The clinical presentations of CL lesions can vary and are capable of self-healing in some cases. However, resolving these can take several months to years at a time, leaving behind a significant amount of scarring. In cases of Post-Kala Azar dermal leishmaniasis, patients can continue to serve as a reservoir for the parasites after the lesions have long been healed [37]. Our results showed significant perturbations in the metabolism of the skin lesions, with the area near the skin lesions also being affected in experimental CL. Our study relied on bioluminescence to measure parasite burden and as such we only have low spatial resolution and cannot ascertain whether parasites were still present at low levels in the sites adjacent to the skin lesions. There is therefore still a strong need to understand the role of lesion-free tissues in transmission of *Leishmania* and in disease pathogenesis.

This study looked at both ear and footpad infection models, although the effect of infection was found to be more minor in the footpad. This could be attributed to the limited sample size for the footpad sampling and a reduced magnitude of metabolic changes, making the study underpowered. Nevertheless, specific PC family metabolites were increased with infection in the footpad, drawing parallels to the ear data and showcasing similarities in pathogenesis processes between these two infection models.

While this untargeted metabolomics study enabled us to uncover several metabolic pathways affected in CL, on average unannotatable compounds (level 2 annotations according to metabolites standards initiative [38]) still represent 88.3% of our data. Molecular networking did enable us to extend annotations further, so that 61.7% of our top 15 most differential metabolite features identified by random forest had at least family-level (level 3) annotations [38]. Nevertheless, metabolomics annotation rates are continuously improving. Our results have been deposited in a “living data” database [23], where they are continuously being re-annotated as reference libraries and computational tools expand. As such, they will continue to yield more insights into CL pathogenesis and serve as a building point for expanded studies of metabolism in CL. Such results will help guide the next generation of CL drug treatments.

## Acknowledgments

Luciferase-expressing *L. major* parasites were provided by Dr. Martin Olivier, McGill University.

## Financial Disclosure Statement

Data collection was supported by a postdoctoral fellowship from the Canadian Institutes of Health Research, award number 338511 to L-IM (www.cihr-irsc.gc.ca/). Work in the McCall laboratory at the University of Oklahoma is supported by start-up funds from the University of Oklahoma (http://www.ou.edu/). This work was also partially supported by US National Institutes of Health (NIH) grant 5P41GM103484-07 to PCD (www.nih.gov/). We further acknowledge NIH Grant GMS10RR029121 (www.nih.gov/) and Bruker (www.bruker.com/) for the shared instrumentation infrastructure that enabled this work (to PCD). The funders had no role in study design, data collection and analysis, decision to publish, or preparation of the manuscript.

## Notes

### Competing Interest Statement

The authors have declared no competing interest.

