## Supplemental Figures for "Metabolite profiling of experimental cutaneous leishmaniasis lesions demonstrates significant perturbations in tissue phospholipids"

### Supporting information

**S1 Fig. PCoA analysis of footpad samples.** (A) PCoA analysis of aqueous extraction from infected (red) and uninfected (blue) footpad samples. PERMANOVA p= 0.244, $R^2$=0.146. (B) PCoA analysis of organic extraction from infected (red) and uninfected (blue) footpad samples. PERMANOVA p= 0.218, $R^2$=0.156.

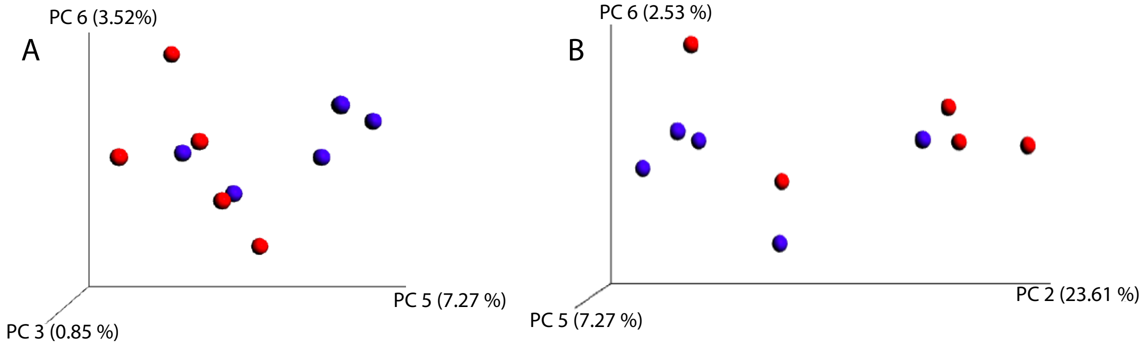

**S2 Fig. Mirror plots for differential annotatable metabolites.** (A) *m/z* 746.6052, RT 4.78, 1-Hexadecyl-2-(9Z-octadecenoyl)-sn-glycero-3-phosphocholine (B) *m/z* 147.0815, RT 0.34, Glutamine (C) *m/z* 792.5574, RT 5.71, Docosaheaenoyl PAF C-16 (D) *m/z* 806.5682, RT 5.42, 1-Palmitoyl-2-docosaheaenoyl-sn-glycero-3-phosphocholine (E) *m/z* 813.6845, RT 5.27, N-Tetracosenoyl-4-sphingenyl-1-O-phosphorylcholine (F) *m/z* 508.3764, RT 4.01, 1-(1Z-Octadecenyl)-sn-glycero-3-phosphocholine (G) *m/z* 522.2834, RT 4.16, 1-(9Z-Octadecenoyl)-sn-glycero-3-phosphocholine (H) *m/z* 303.2323, RT 4.1, 5,6-Epoy-8Z,11Z,14Z-eicosatrienoic acid (I) *m/z* 744.5891, RT 6.01, 1,2-Di-(9Z-octadecenoyl)-sn-glycero-3-phosphoethanolamine (J) *m/z* 794.6035, RT 5.97, (2-{[2-[icosa-5,8,11,14-tetraenoyloxy]-3-[octadec-1-en-1-yloxy]propyl phosphonato]oxy}ethyl)trimethylazanium (K) *m/z* 796.6135, RT 6.64, 1-Heptadecanoyl-2-(5Z,8Z,11Z,14Z-eicosatetraenoyl)-sn-glycero-3-phosphocholine (L) *m/z* 813.6867, RT 7.51, N-Tetracosenoyl-4-sphingenyl-1-O-phosphorylcholine.

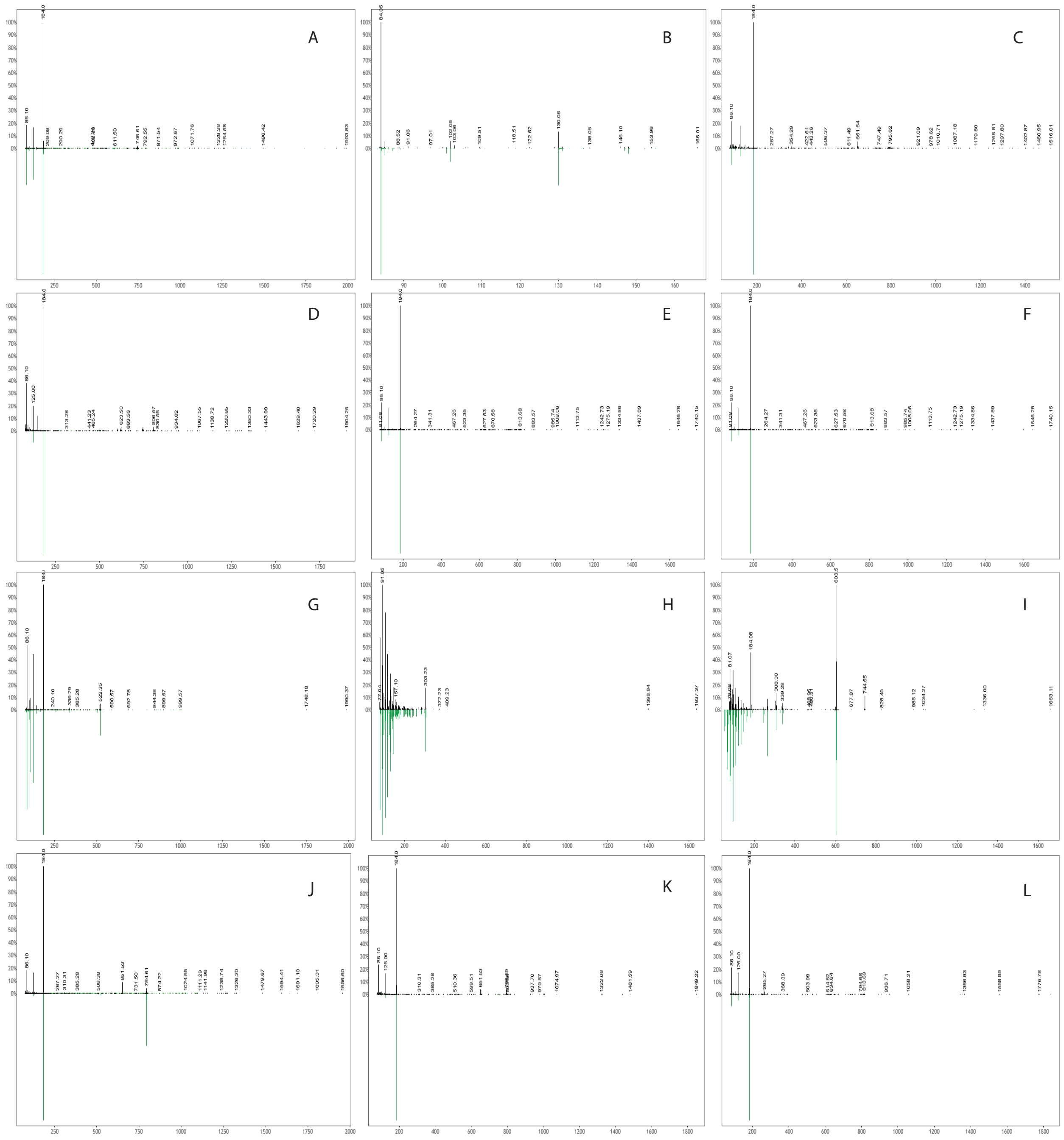
A
B
C
D
E
F
G
H
I
J
K
L

**S3 Fig. PC subnetworks.** (A-D) PC family metabolites in the ear aqueous (A), ear organic (B), footpad aqueous (C) and footpad organic molecular networks (D). Metabolite abundance in the presence of infection is shown in red and absence of infection is blue. Node size is directly proportional to peak area (individual scales for each sub-panel).

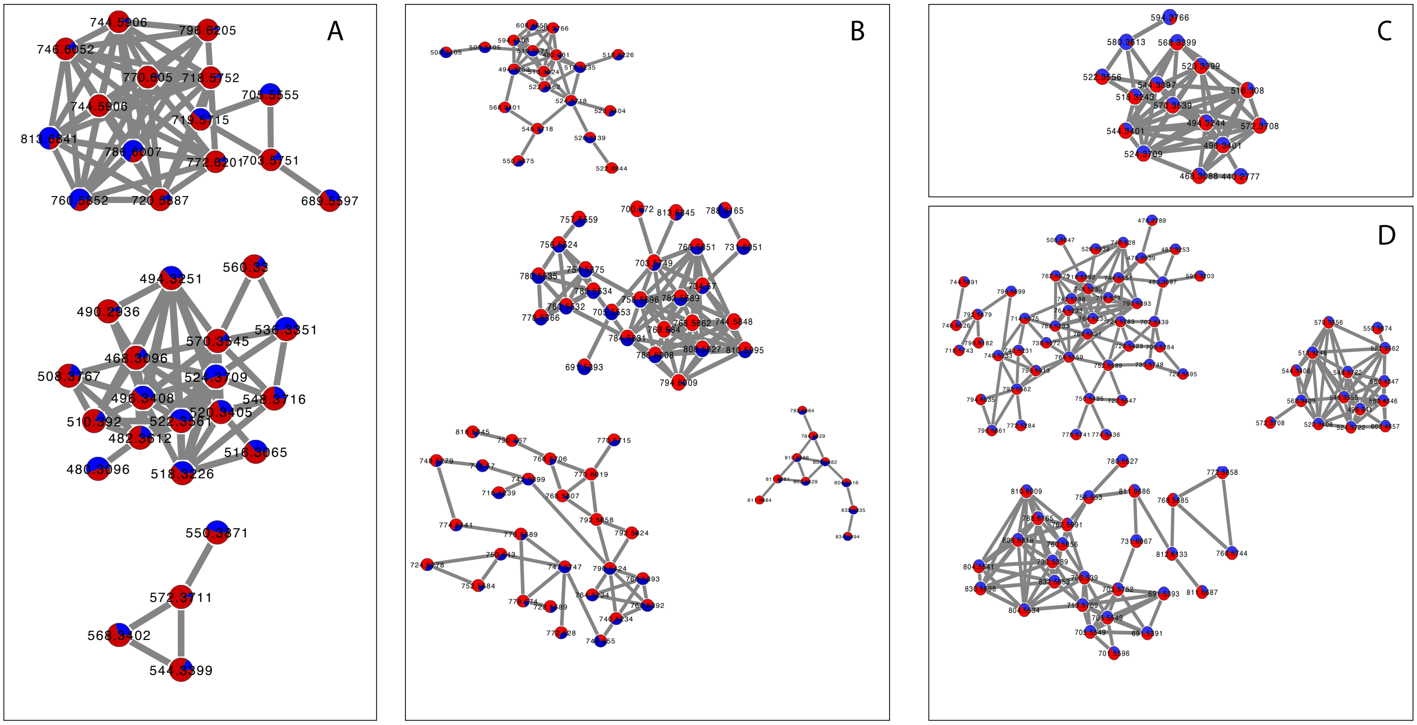
